# The impact of *Cronartium ribicola* inoculum density on quantitative disease resistance in whitebark pine

**DOI:** 10.64898/2026.05.02.722345

**Authors:** Jeremy S. Johnson, Benjamin Wilhite, Robert Danchok, Angelia Kegley, Richard A. Sniezko

## Abstract

Whitebark pine (*Pinus albicaulis*), a wide-ranging high-elevation conifer in western North America, is listed as threatened in the U.S. and as endangered in Canada. A major threat to whitebark pine is the non-native, invasive white pine blister rust disease, caused by the fungal pathogen *Cronartium ribicola*. In many pathosystems (including white pine blister rust), seedling inoculation trials are used to identify parent trees with genetic resistance. However, many of these trials use only one spore density for inoculation, and little information exists on the effectiveness of quantitative disease resistance (QDR) under varying spore densities and the corresponding implications for field performance. In this study, we examine the levels of infection and survival present within six whitebark pine seedling families previously rated for QDR (three susceptible and three resistant families) under six widely varying inoculum densities. The susceptible families showed very high infection and mortality at all inoculum densities, while performance of the resistant families varied with spore density treatment. The information gathered from the study will be useful in updating the projections of the future of whitebark pine populations under field conditions in areas of different rust hazard. The results also serve as a caution to those working in other pathosystems where seedling inoculation trials based on one spore density level are used to rate the resistance level of parent trees and their associated progeny.

## Introduction

*Pinus albicaulis* Engelm. (whitebark pine) is a long-lived conifer and keystone species that grows in high elevation forests throughout the western U.S. and Canada (Tomback et al. 2022). It is one of nine species of five-needle or white pines in the subgenus *Strobus* found in the U.S., four of which are also found in Canada. All North American species of five-needle pines are highly susceptible to white pine blister rust (WPBR), caused by the non-native, invasive pathogen *Cronartium ribicola* J.C. Fisch. in Rabh. (Hoff et al. 1980, Tomback and Achuff 2010). Because *P. albicaulis* is impacted by multiple threats, including high mortality from WPBR, predation from mountain pine beetle (*Dendroctonous ponderosae* Hopkins), changes in fire regimes and the climate (Tomback et al. 2022), it has been listed as endangered in Canada under the Species at Risk Act (SARA) in 2012 (Government of Canada, 2012) as well as listed as a threatened species in the U.S. under the Endangered Species Act (ESA) (U.S. Fish and Wildlife, 2022). *P. albicaulis* is also listed as endangered on the IUCN Red List (Mahalovich and Stritch 2013). In the U.S. a national plan for restoration of whitebark pine has been proposed (Tomback and Sprague 2022). Identifying trees with genetic resistance to WPBR is vital to the success of restoration (Schwandt 2006) to keep white pine species as functioning components of North American forests.

White pine blister rust is a serious disease of five-needle pine trees in North America where it has been present for over 100 years and is now considered a permanent inhabitant of many white pine ecosystems (Kinloch 2003, Geils et al. 2010). *Cronartium ribicola* is an obligate, biotrophic pathogen that requires both a telial host (usually *Ribes* species in North America) and an aecial host (white pine species including *P. albicaulis*). Infection in the pines occurs when basidiospores are dispersed from the telial host and enter through the needle stomata (Geils et al. 2010). As the pine becomes infected, the fungus will grow from the needle tissue into the stem where one or more cankers may develop (Geils et al. 2010). Both the needles and stems of the pines are potential areas where resistance responses restricting progress of the pathogen may occur. In the stem, this may be apparent by slower growing cankers or partial or complete bark reactions that disrupt the life cycle of the fungus (Hoff 1986, 1992).

Fortunately, native white pine species have some natural genetic resistance to the non-native white pine blister rust (Kinloch et al. 1970, Hoff and McDonald 1980, Sniezko et al. 2014, Sniezko and Liu 2022). Though resistance mechanisms and their effectiveness vary, two types of host resistance are typically recognized—major gene resistance (MGR) and quantitative disease resistance (QDR) (Liu et al. 2021). MGR is a low frequency race-specific resistance, conditioned by a single major gene, that can completely arrest the spread of the fungus through a hypersensitive-like reaction (HR) in the needles (Kinloch and Dupper 2002). The formation of necrotic needle lesions (spots) disrupts the ability of the fungus to access the living host tissue which it needs to spread. Alternatively, QDR is a non-race-specific form of resistance, influenced by many genes, that reduces the impacts of the disease, and in some cases results in the complete absence of the disease in stems (Johnson and Sniezko 2021, Liu et al. 2021). QDR is found at low frequencies in many white pine species including *P. albicaulis*, but to date MGR has not been documented in whitebark pine (Sniezko et al. 2024). Recent and extensive investigations of *P. albicaulis* resistance through artificial seedling inoculation trials have provided some optimism concerning the frequency and level of QDR to white pine blister rust in this species (Hoff et al. 2001, Sniezko et al. 2007, Sniezko et al. 2008, Sniezko et al. 2011, Sniezko and Liu 2022, Sniezko et al. 2024). Compared to other white pine species in the U.S., open-pollinated seedling progeny of resistant *P. albicaulis* parent trees have a higher, more usable level of QDR without breeding, with survival frequencies of 40% or greater in some progenies (Sniezko et al. 2024). However, since most seedling inoculation trials testing for WPBR resistance utilize only one spore density (generally selected to ensure a robust test and minimizing escapes), this could result in under or overestimation of the level of resistance, particularly under field conditions.

Durability of resistance is a concern in all plant pathosystems but is paramount for forest tree species, where trees are expected to be on the landscape for decades to centuries after reforestation or restoration activities (Sniezko and Liu 2023). In the case of restoration, the expectation is that the resistance will remain effective over generations and provide a new equilibrium where the host species can continue to exist and provide ecosystem services even in the continued presence of the pathogen. Evolution of the pathogen towards greater virulence can occur, which can be a major impediment to durability of MGR, but QDR is generally expected to be more durable than MGR (Sniezko and Liu 2023). Change in virulence of the pathogen is not the only factor that may influence the utility of resistance. As shown in WPBR field trials and field surveys with *P. monticola* and *P. lambertiana*, environments can vary greatly in disease hazard (Koester et al. 2018, Sniezko et al. 2020), and much less is known (few or no reports in the literature) about how disease hazard, defined as the density of spores at a particular site with a suitable environment for infection, can affect survival of populations with QDR. If the efficacy of QDR in whitebark pine varies radically under increasing disease pressure, then the distribution of environments with low to extreme disease hazard will ultimately influence the utility of QDR of a species (Sniezko et al. 2020) and the success of natural regeneration or restoration plantings.

Several open questions remain regarding how site-level hazard of *C. ribicola* basidiospores varies among environments and how this variation impacts the effectiveness of QDR expressed in the progeny of maternal trees. In particular, it is unclear how increasing inoculum pressure changes survival and disease severity across resistant and susceptible families. To address these questions, we developed an artificial seedling inoculation experiment to test the overarching hypothesis that the effectiveness of QDR in *P. albicaulis* is context dependent and declines as *C. ribicola* inoculum pressure increases, such that increasing basidiospore density increases mortality risk and disease severity across all families, which will reduce, but not eliminate, the survival advantage of QDR families compared to susceptible families. The results presented may suggest caution for resistance programs in other pathosystems to avoid potential over-estimation or under-estimation of resistance levels from artificial inoculation trials.

## Materials and Methods

### Experimental design

Open-pollinated seeds of six whitebark pine maternal trees (families) were stratified in fall 2017, sown in Spring 2018, and grown in an unheated greenhouse for two growing seasons. Two-year old seedlings of the six families were inoculated in September 2019 (Fig. 1a). Seedlings within a family were divided into six treatment groups to be inoculated by increasing levels of spore density. Seedlings were in tubes when inoculated with *C*. *ribicola* then transplanted into 0.9 m x 1.2 m x 0.3 m boxes outdoors for the remainder of the experiment (Fig. 1b).

**Figure 1.**
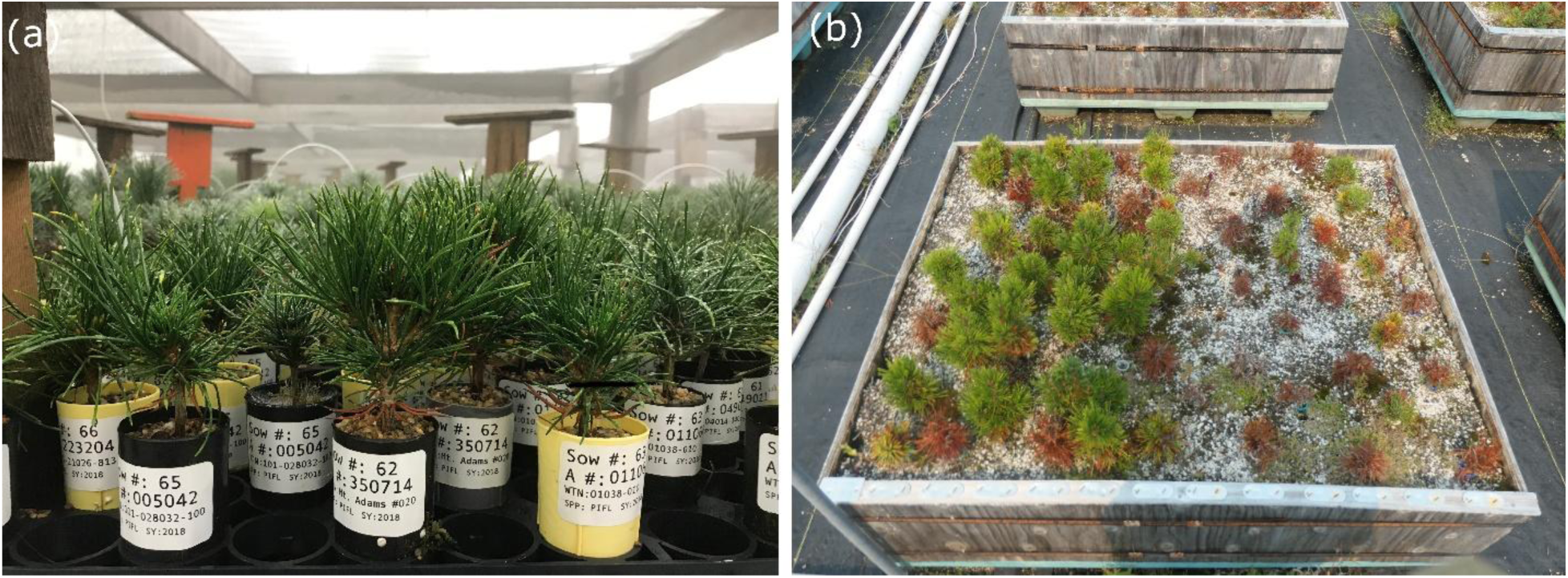
Whitebark pine seedlings from the Sow Year 2018 spore density trial at Dorena Genetic Resource Center (a) in the inoculation fog chamber in September 2019, and (b) in a box outdoors in August 2021 showing high contrast in survival between resistant family sow 66 (left six rows) and susceptible family sow 65 (right six rows) for the six spore density treatments.

Based on prior resistance testing and availability of seed, we selected three high QDR resistant (A) and three of the most susceptible (F) families (Table 1). After inoculation, seedlings were planted in family rows within the wooden boxes, where all seedlings in a row belonged to one family by treatment combination. Each box contained a total of 12 rows, with six assigned to a resistant family, and the other six assigned to a susceptible family (Fig. 2). Rows within box were grouped by family, and treatment was randomized within that grouping. The number of seedlings planted per row varied from three to seven (mean = 6). A total of nine boxes were used, with each family being present in three boxes (each A family paired with a different F family in each of the boxes) (Fig. 2).

**Figure 2.**
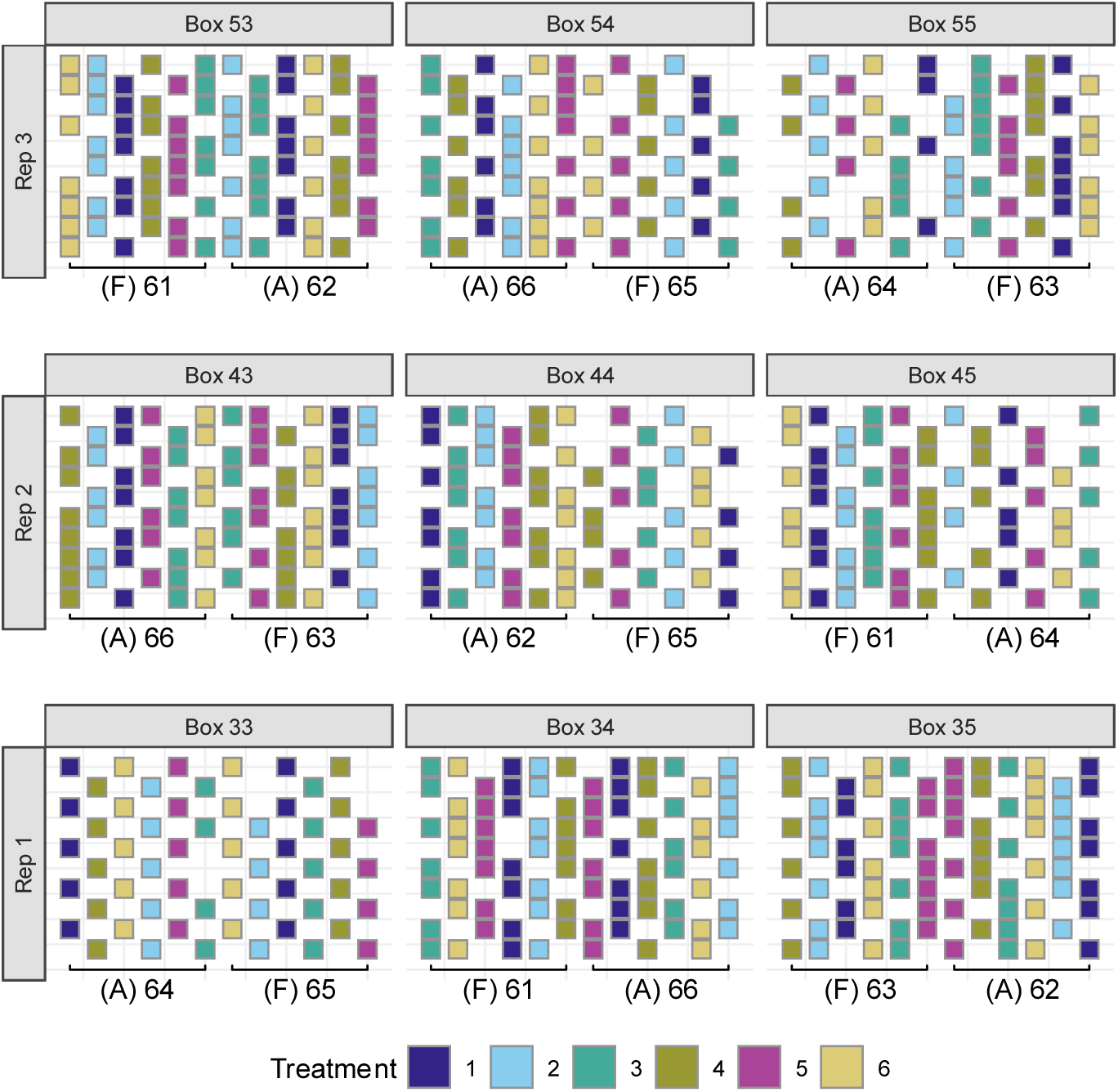
Whitebark pine seedling map illustrating the study design. Seedlings were arranged in family by treatment row-plots, with each box including all family by treatment combinations for two families, one resistant and one susceptible. Brackets indicate the resistance level (A = resistant, F = susceptible) and Sow number of each family. Each replicate consisted of three boxes that, together, included every family by treatment combination.

**Table 1.** Identities of the six whitebark pine families, prior blister rust resistance rating, rating year, and number of seedlings in each of the six spore density treatments.

| Sow<br>No. | Seed Source | Year<br>Rated | Rating | No.<br>Reps | No. Seedlings Per Treatment |  |  |  |  |  |  |
| --- | --- | --- | --- | --- | --- | --- | --- | --- | --- | --- | --- |
|  |  |  |  |  | T1 | T2 | T3 | T4 | T5 | T6 | Total |
| 62 | Mt Adams #020 | 2012 | A | 3 | 21 | 21 | 21 | 21 | 21 | 20 | 125 |
| 64 | CL18 | 2007 | A | 3 | 14 | 14 | 14 | 14 | 14 | 13 | 83 |
| 66 | 101-21026-813 | 2007 | A | 3 | 20 | 20 | 20 | 20 | 20 | 19 | 119 |
| 61 | 04014-140 | 2004 | F | 3 | 21 | 21 | 21 | 21 | 21 | 17 | 122 |
| 63 | 01038-010 | 2007 | F | 3 | 19 | 20 | 20 | 20 | 20 | 20 | 119 |
| 65 | 101-028032-100 | 2013 | F | 3 | 14 | 14 | 14 | 14 | 14 | 13 | 83 |

### Inoculation

Artificial inoculation followed standard procedures (Kegley and Sniezko 2004). In brief, seedlings were inoculated in a fog chamber, which was held at a constant 16.6°C and 100% relative humidity. Infected *Ribes* leaves with visible *C. ribicola* telia were placed telia side down on wire mesh frames above the seedlings (Fig. 1a). Seedlings were placed into the inoculation fog chamber on September 16, 2019, and removed on September 23, 2019. Spore fall occurred September 19, 2019, and leaves were removed on September 20, 2019. Spore fall was monitored by counting spores on adhesive covered glass slides that were dispersed among the seedlings. Once the desired inoculum densities were achieved, the infected *Ribes* leaves were removed from the screens above the seedlings. Treatments one through six averaged 593, 1367, 3947, 7653, 15020, and 24500 spores cm^-2^, respectively. Treatment three (3,947 spores cm^-2^) was the closest to the standard inoculation density used at the USDA Dorena Genetic Resource Center (DGRC) (∼3000 spores cm^-2^). The lowest density (treatment one) is approximately 20% of DGRC standard, and the highest density (treatment six) is approximately 800% of DGRC standard levels.

### Tree Inspection

For the five-year duration of the trial, all seedlings were periodically monitored for WPBR disease symptom development. Disease symptom inspections began in May 2020, approximately 228 days post-inoculation (dpi), and continued for 1,959 days. At each inspection, seedlings were assessed for disease symptoms, severity, and survival. Severity of WPBR infections was visually assessed and quantified on a 0 to 9 scale, where 0 = no stem symptoms, 1 to 4 = one or more stem symptoms with only minor overall impact (canker <100% stem diameter), 5 to 8 = one or more cankers encircling 100% of the stem and increasing vertical growth of the cankers for categories 6 to 8, with seedlings ranked as 8 having nearly the entire stem cankered, and 9 = dead from WPBR.

Primary needle spots, the first observable disease symptoms, were counted up to 50, and then in blocks of 25, at the first inspection (228 dpi) and then counted on secondary needles up to 50, and then in blocks of 25, at the second inspection (458 dpi). Spots were also recorded as a binary variable (presence or absence) at both inspection. Subsequent inspections did not evaluate the presence of needle spots. Number of stem symptoms on the bole (main stem of the seedling) and total number of stem symptoms occurring on either the bole or branches of each seedling were counted at second inspection (458 dpi). In this trial, some merging of stem symptoms was present at second assessment, particularly in the higher spore density treatments, and the counts may have underestimated the actual number of cankers present. Based on this, for inspections three through six, the presence of stem symptoms was recorded as a binary measure, and the composite stem symptom phenotype (stem symptoms observed on bole or branches) was treated as a presence or absence variable for analysis.

## Statistical Analysis

### Survival Analysis

Survival was summarized as time to rust associated death over the six assessments conducted during the five-year period following inoculation. Observations were mapped to dpi. For each tree, survival status was coded as zero if the seedling was first recorded dead at any assessment and 1 if it remained alive through the final assessment. Survival time was defined as the first inspection time at which mortality was observed, or the final observed inspection time where death was not observed (censored seedlings). Trees that died from causes other than WPBR were excluded from the analysis.

### Cox Regression Time to Mortality

In order to evaluate how family level survival through time was impacted by different levels of inoculum density, we used a Cox proportional hazards regression model. The model estimated the effect of treatment on the hazard of white pine blister rust mortality. Because open-pollinated seedlings come from the same maternal family, they are not statistically independent. We calculated standard errors by clustering observations at the family level. The first model tested if inoculum density had an impact on the risk of mortality at any given time following inoculation, where survival time is defined as the number of days post inoculation until mortality was first observed or the final inspection data (Eq1).

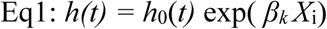

where *h*(*t*) = hazard of mortality at time *t, h*_0_(*t*) = baseline hazard at treatment one, *X_i_* = inoculum density at treatment level, and *β_k_ =* estimated effect of treatment on mortality. Hazard ratios are: *HR* = exp(*β_k_*); values of HR > 1 indicate increased mortality risk.

A second Cox model (Eq. 2) was fitted to evaluate if the effect of family genetic resistance rating (resistant vs. susceptible) on survival varied across inoculum densities. This model included the family resistance rating, inoculum density treatment, and their interaction. The interaction term tested whether resistant families retained a survival advantage with increasing spore density and whether the magnitude of the advantage changed across treatments. Kaplan–Meier step-function curves were used to visually assess patterns of survival.

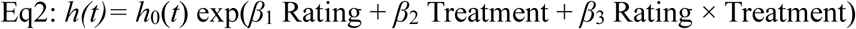

where *h*(*t*) = hazard of mortality at time *t, h*_0_(*t*) = baseline hazard at treatment one, Rating = resistance score (A = resistant and F = susceptible), Treatment = inoculum density level relative to the baseline treatment one, and Rating x Treatment represents their interaction.

Hazard ratios are: *HR* = exp(*β*); values of HR > 1 indicate increased mortality risk relative to the baseline, and HR < 1 indicates reduced mortality risk.

### Repeated Measure Analysis

Because we assessed survival at discrete inspections, we also analyzed the probability that a seedling would be alive at each inspection using generalized linear mixed-effects models with a binomial error distribution and logit link (Eq3). We included treatment, inspection, and their interaction as fixed effects, with family included as a random intercept to account for non-independence among seedlings from the same maternal family. A second model assessed resistance rating (A or F) and included a rating by treatment interaction to evaluate if resistant and susceptible families differed in temporal patterns of survival across inoculum densities.

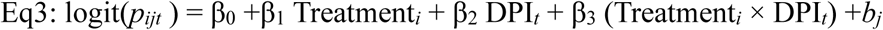

Where: the probability that seedling *i* in family *j* is alive at days post inoculation *t.* logit(*p_ijt_*) = the log odds of being alive, β_0_ = intercept, Treatment = inoculum density level, DPI = days post inoculation, Treatment × Inspection = if density effects change through time, *b_j_* = random intercept for family and *b_j_* ∼N(0,σ^2^_family_)

### Needle Spot and Canker Counts

Needle spot and normal canker counts were analyzed using generalized linear mixed effects models. We fitted a negative binomial mixed-effects model (Eq4) with treatment as a fixed effect and family as a random effect. The initial model was compared to a zero-inflation negative binomial model using AIC.

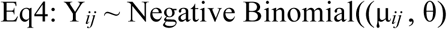

Where: Y*_ij_ =* observed needle spot count, μ*_ij_* = expected needle spot count for seedling *i* in family *j*, θ = negative binomial dispersion parameter, β_0_ = intercept for the expected log spot count of treatment one, β_k_ = fixed effect coefficient for treatment *k* relative to baseline treatment one, Treatment*_ik_* = variable equal to 1 if seedling *i* was assigned to treatment *k* and 0 otherwise, *b_j_* = random intercept for family *j,*,σ^2^_family_ = variance among families in baseline treatment one.

Residual diagnostics, including tests of dispersion and zero inflation, were evaluated using the DHARMa package in R (Hartig 2026). Although residual diagnostics suggested a deficit of zeros, the zero<u>-</u> inflation model did not improve the fit based on AIC (negative binomial = 4927; zero negative binomial = 4929) and was not retained for further analysis. Treatment was analyzed first as a six-level factor to estimate treatment-specific effects on needle spot counts. When treatment effects were significant, estimated marginal means and Tukey-adjusted pairwise comparisons were used to compare the impact of inoculum density on the number of needle spots. Similarly, normal canker counts were analyzed at inspection two (458 dpi) prior to significant canker merging at later inspections.

### Dose-response of needle spot severity and canker count

To test whether disease severity, needle spots, and canker counts increased linearly or nonlinearly with increasing inoculum density, spore density was analyzed as a continuous predictor following log_10_ transformation to account for the wide range of densities. Model predictions were then back transformed to the original scale (spores cm^-2^) for interpretation and visualization on a log_10_ axis. Negative binomial mixed-effects models were fitted with family as a random effect.

Models were fitted with either a linear term of log_10_ spore density only or both linear and quadratic terms. The models were compared using likelihood ratio tests and AIC. We interpreted a better fit of the quadratic model as evidence of a nonlinear dose dependent relationship with saturation of disease severity occurring at higher inoculum density. We calculated the point at which increasing spore density resulted in no further increase in disease symptoms based on the best fit model (Eq5).

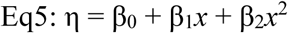

Where η is the linear predictor, *x* = log_10_ (spore density), and β_0._ β_1, and_ β_2_ are model coefficients.

The turning point of the quadratic function is 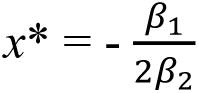

### Survival dose-response analysis

Seedling survival was analyzed using binomial generalized linear mixed-effects models that included family as a random effect, and log_10_ spore density and dpi were included as fixed effects. Alternate models were compared to test for nonlinear dose response and the interaction between resistance rating and spore density. AIC and likelihood ratio tests were used to select the best model. Odds ratios were calculated to quantify the changes in survival probability associated with a ten-fold increase in spore density.

### Resistant family differences in survival

Lastly, we explored whether families previously tested and rated A and thus QDR resistant differed in their patterns of survival in response to increasing spore density. Scaled spore density and inspection time were modeled in a binomial mixed model, and a random-intercept model was compared with a random-slope model allowing the effect of spore density to vary by family. Model support was evaluated using likelihood ratio tests and AIC.

Unless otherwise noted, statistical significance was evaluated at 95% confidence (α = 0.05). Descriptive plots of percent survival by family, treatment, and inspection are used to illustrate the biological pattern, whereas our inferences are based on the models described above. All analysis was carried out within the R statistical environment (RCoreTeam 2026).

## Results

Inoculation was very effective with nearly every seedling developing needle spots at first assessment (99.1%), and all but one seedling showed spots at first or second assessment and/or stem symptoms. There was some variability within families and between treatments with mean percent needle spots at 228 dpi ranging between 90.5 and 100%. High mortality (family by treatment range = 0-100%; µ = 64%) from rust infection was apparent within three and half years of inoculation (1,259 dpi).

### Survival Analysis

Percent survival over time varied widely among families and across treatments. All families and treatments had 100% survival at 458 dpi. By the end of the experiment, survival ranged from 0 to 93% for treatment one and 0 to 35% for treatment six across families (Fig. 3). The overall mean survival across treatments and families was 14%.

**Figure 3.**
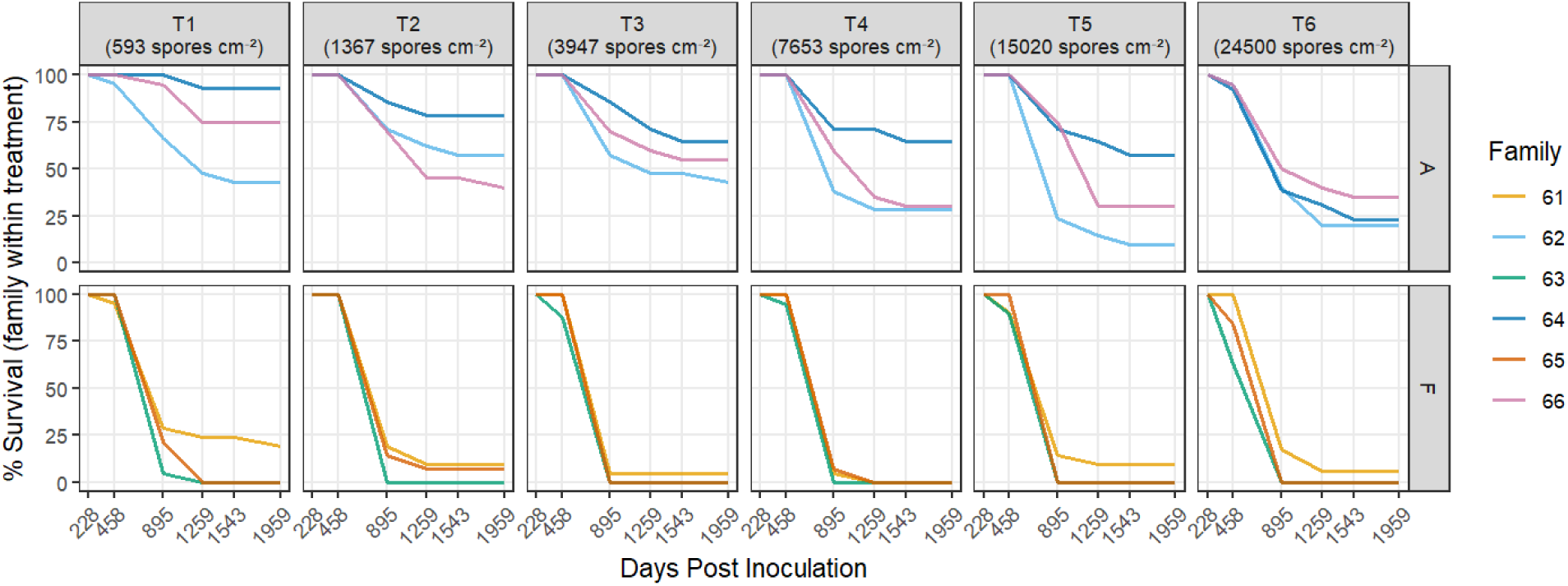
Percent survival of whitebark pine by family and spore density treatment over time. Lines represent family percent survival across treatments (colors correspond to family in the legend, and treatment and spore density are labeled across the top of the panels). Panel represents resistance rating A (top) = resistant and F (bottom) = susceptible. The x axis shows the number of days post inoculation (dpi).

### Cox Regression Time to Mortality

A total of 653 seedlings were assessed with 494 mortality events observed during the experiment. Time to rust associated death over six assessments was analyzed using a Cox proportional hazard regression model. Treatment significantly affected the probability of mortality (Wald test, *p* <0.001). Relative to a baseline at treatment one, mortality risk increased with increasing spore density (Table 2). Seedlings in treatment three had a 37% greater risk of mortality (hazard ratio = 1.37, 95% CI: 1.15-1.65), with treatments four and five increasing mortality risk by 67% (HR = 1.67, 95% CI: 1.25-2.22) and 76% (HR = 1.76, 95% CI: 1.20-2.37), respectively. The greatest risk of mortality occurred in treatment six, where seedlings had about twice the risk of mortality compared to treatment one (HR=2.01, 95% CI: 1.29-2.89). Treatment two did not differ significantly from treatment one (HR = 1.2, 95% CI: 0.87-1.67, *p* = 0.26). The model accuracy was 0.61, which indicated that it was able to differentiate survival outcomes about 61% of the time. There were no violations of proportional hazard assumptions for the model based on Schoenfeld residuals (global test, *p* = 0.7).

**Table 2.** Cox proportional hazard model. The model indicates a significant difference in mortality between treatments with an increase in whitebark pine mortality associated with an increase in spore density (spores cm^-2^). Hazard Rating (HR) is the expected increase in probability of mortality assessed at 95% confidence (CI). Bolded *p-*values are significantly different from the treatment one base.

| Treatment | Spore Density<br>(spores cm<sup>-2</sup>) | HR<br>(exp(coef)) | 95% CI | *p*-value |
| --- | --- | --- | --- | --- |
| 1 | 593 | - | - | - |
| 2 | 1,367 | 1.2 | (0.87–1.65) | 0.26 |
| 3 | 3,947 | 1.37 | (1.15–1.65) | **0.0006** |
| 4 | 7,653 | 1.67 | (1.25–2.22) | **0.0004** |
| 5 | 15,020 | 1.76 | (1.30–2.37) | **0.0002** |
| 6 | 24,500 | 2.01 | (1.39–2.89) | **0.00018** |

A mixed-effects logistic regression model tested the probability of survival across inspections with treatment, dpi and their interactions as fixed effects and family as a random intercept. The model found strong time-based effects with probability of survival decreasing significantly with dpi (*X^2^* = 183.90, df = 1, *p* <0.001). Treatment at 228 dpi was not a significant predictor of survival (*X^2^* = 1.77, df = 5, *p* =0.88). The treatment by dpi interaction was significant (*X^2^*= 33.48, df = 5, *p* < 0.001), suggesting that the rate of survival decline over time differed among treatments. Family level heterogeneity in survival was observed (random effect SD = 1.45). Both the family level and aggregated models showed signs of dose-dependent mortality.

Another mixed-effects logistic regression tested the probability of tree survival across six inspection periods as a function of treatment (spore density), with a random intercept for family. Family explained substantial variation in survival (SD = 2.0), indicating strong genetic differences in resistance. Survival declined sharply after the second inspection in all treatments, but higher inoculum doses (treatments three through six) showed significantly larger declines through time, as reflected by strong negative treatment by inspection interaction terms.

Treatments one and two showed similar temporal survival patterns. Because there was nearly 100 percent mortality at later inspections, several logistic coefficients were large, and the model showed signs of almost-complete separation between families. However, this is not uncommon in cases with high rates of mortality. The overall pattern and separation illustrate a dose-dependent acceleration of mortality and strong family-level genetic effects of QDR.

Kaplan-Meier survival curves showed large differences in survival between resistant (A rating) and susceptible (F rating) families following inoculation (Fig. 4). Survival of susceptible families declined, beginning at 895 dpi. On balance, resistant families maintained a higher survival advantage for the duration of the experiment. At the end of the experiment (1,959 dpi), resistant families had close to 45% survival while susceptible families were at 10%. The log-rank test indicated a statistically significant difference in survival between resistant and susceptible families (*p* < 0.001).

**Figure 4.**
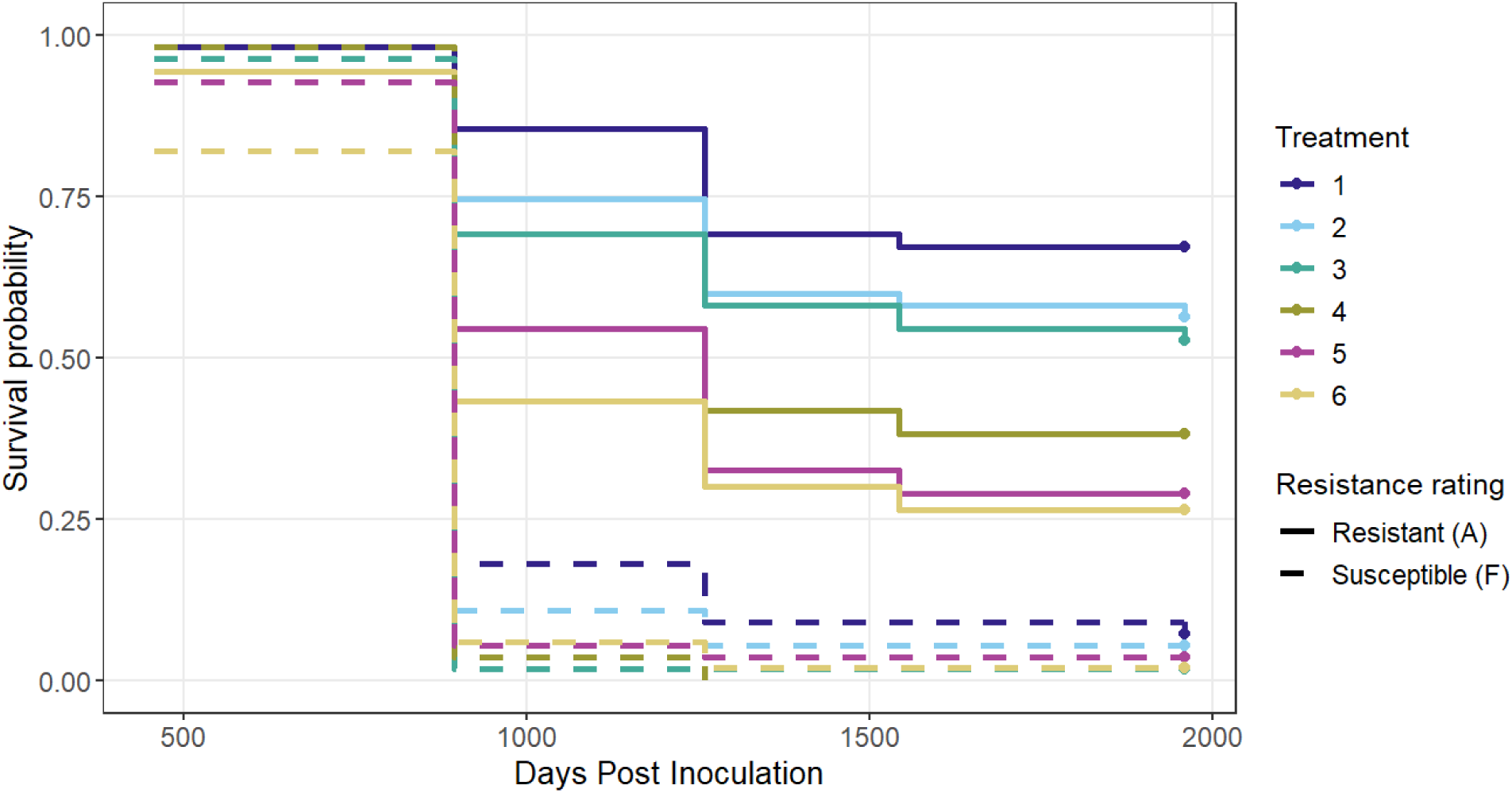
Kaplan-Meier survival curves for whitebark pine seedlings grouped by resistance rating and treatment (inoculum density). Solid lines are resistant families (rated A), and dashed lines are susceptible families (rated F). Line color indicates treatment and corresponding inoculum density (Table 1). Survival declined with increasing inoculum density. Moreover, inoculum density reduced the overall survival advantage of families with resistance to white pine blister rust relative to susceptible families.

In addition to the Kaplan-Meier analysis, a second Cox proportional hazard model was used to evaluate the effect of resistance rating, inoculation density treatment, and their interaction on mortality risk. Resistance rating significantly influenced probability of mortality, with susceptible families possessing almost seven times greater risk of mortality compared with resistant families (HR = 6.95, 95% CI: 2.53-19.12, *p* <0.001). An increase in spore density also resulted in increased mortality among resistant families. Compared to treatment one, mortality risk increased in treatment four (HR=2.58, *p* < 0.001), treatment five (HR = 3.07, *p* <0.001), and treatment 6 (HR = 3.66), p < 0.001) (Table 3). A significant resistance rating by treatment interaction suggested that the strength of the treatment effect differed between resistant and susceptible families. This was particularly apparent at the highest inoculum densities (treatment five: HR = 0.46, *p* < 0.001 and treatment six: HR = 0.44, *p* = 0.040). In combination, the results showed that even though resistant families had much greater overall percent survival, increasing spore density still reduced survival across resistance classes, and particularly decreased the advantage of resistant families at higher inoculation densities (Fig. 3). The overall model performance was good with approximately 83% of the variation in mortality explained across seedlings.

**Table 3.** Hazard ratio (HR), Wald z-score, and *p –* value from a Cox proportional hazard regression model evaluating the effect of inoculum density treatment on the hazard of white pine blister rust associated mortality. Hazard ratios are expressed relative to treatment one (considered as the reference level). Values of HR > 1 indicate increased risk of mortality compared to the reference.

| **Treatment** | **HR** | **z** | ***p*-value** |
| --- | --- | --- | --- |
| **2** | 1.46 | 0.824 | 0.41 |
| **3** | 1.66 | 1.76 | 0.078 |
| **4** | 2.58 | 3.252 | **<0.001** |
| **5** | 3.07 | 7.197 | **<0.001** |
| **6** | 3.66 | 3.205 | **<0.001** |

### Needle Spot and Canker Counts

White pine blister rust needle spots increased with higher levels of spore density. A negative binomial mixed-effects model showed a sizeable dose-dependent increase in expected spot counts across treatment levels (likelihood ratio test, *p* <0.01). Relative to treatment one, mean spot counts approximately doubled in treatment two and increased nearly ten-fold in treatment six (Table 4).

**Table 4.** Estimated marginal mean whitebark pine needle spot counts and 95% confidence intervals for inoculum density treatments. Estimates were made from a negative binomial mixed effects model. Treatment was included as a fixed effect, and family was a random intercept. Means are shown on a needle spot response scale.

| Treatment | Mean spots | 95% CI |
| --- | --- | --- |
| 1 | 3.69 | 2.64 – 5.15 |
| 2 | 7.39 | 5.33 – 10.25 |
| 3 | 21.79 | 15.79 – 30.07 |
| 4 | 20.50 | 14.84 – 28.31 |
| 5 | 30.33 | 21.99 – 41.81 |
| 6 | 36.70 | 26.56 – 50.73 |

Estimated marginal means increased from 3.7 spots per seedling (95% CI: 2.6-5.2) in treatment one to 36.7 spots per seedling (95% CI: 26.6-50.7) in treatment six. Tukey pairwise comparisons indicated that treatments three and four did not differ significantly, nor did treatments five and six. All other treatment comparisons were significant (*p* < 0.05).

Family identity explained additional but comparatively modest variation in needle spot counts (random intercept SD = 0.36). This indicates some shared genetic or environmental differences in infection intensity among families.

Model diagnostics indicated that the negative binomial distribution appropriately accounted for overdispersion (dispersion = 0.835, *p-*value = 0.804). Residual diagnostics suggested fewer zeros than expected under the model. However, zero inflated models did not improve model fit based on AIC (negative binomial = 4927 and zero negative binomial = 4929) and were not retained. Infection was nearly universal (∼99.1% of seedlings with at least one needle spot), and the small deficit of zero counts relative to model expectations is consistent with high infection success following inoculation.

At 458 dpi, normal canker counts increased nonlinearly with increasing spore density (Fig. 5). Model comparisons identified a quadratic model over a linear response (ΔAIC = 10.1; *X*^2^ = 14.13, df = 2, *p* < 0.001). The strongest increase occurred at low to moderate spore densities and plateaued at the highest density. Canker counts may be partially underestimated at high spore density treatments due to small seedling size and more extensive canker merging making true counts more difficult. Susceptible families had more cankers compared to resistant families across all treatments. Predicted marginal mean normal cankers ranged from 1.82-3.03 cankers per tree in susceptible families compared to 0.61-1.32 in resistant families. This pattern suggests that higher spore hazard increased early stem symptom occurrence in both resistant and susceptible rated families with resistant trees having a relative advantage.

**Figure 5.**
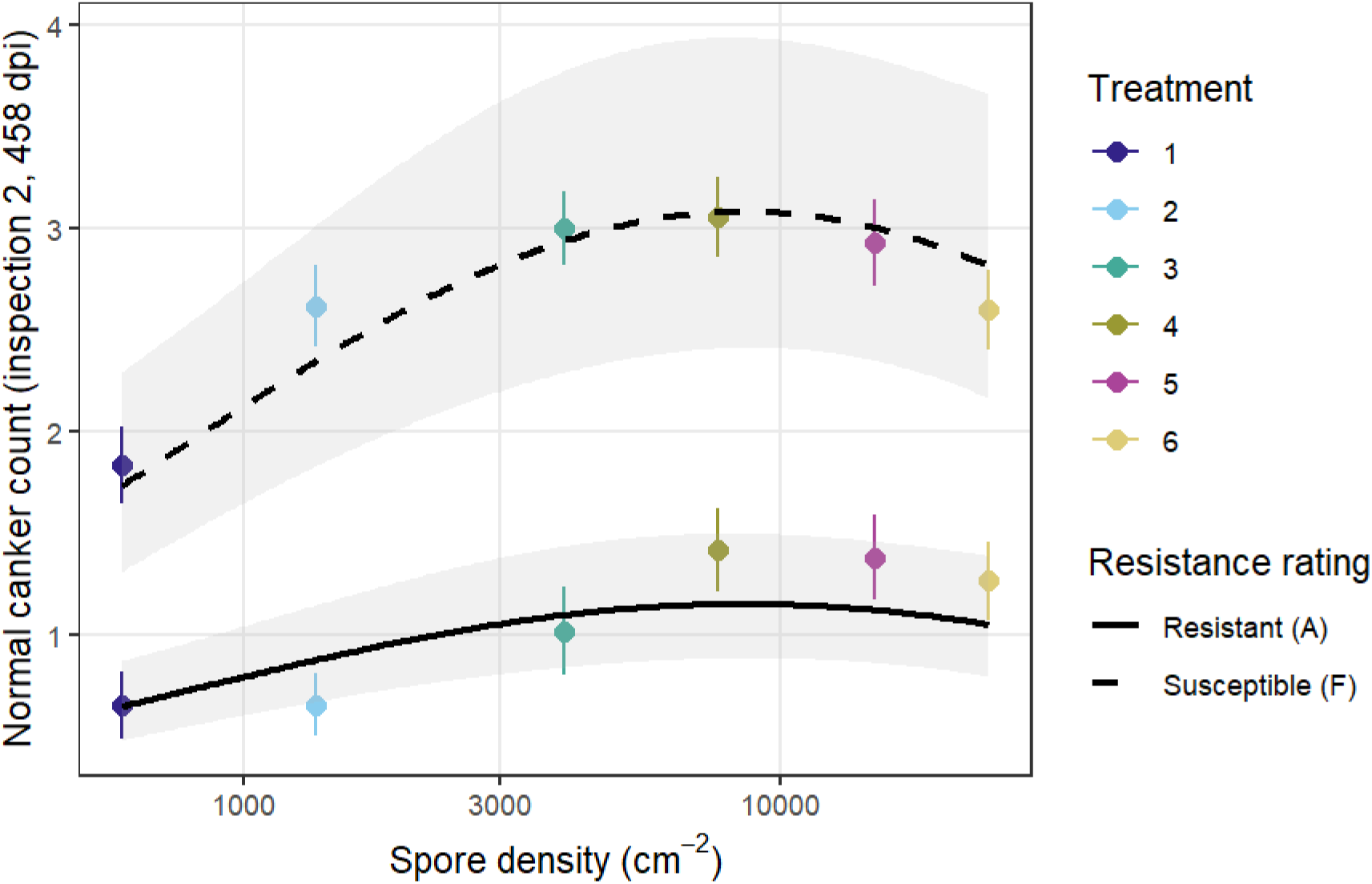
Predicted early stem symptom development (normal cankers) counts at 458 days post inoculation for whitebark pine seedlings as a function of spore density. Points represent observed treatment means (±SE) for resistant (A, solid line) and susceptible (F, dashed line) families.

Model diagnostics indicated that the negative binomial distribution appropriately accounted for overdispersion (dispersion = 0.933, *p* = 0.696) but did find a slight excess of zeros (zero inflation test *P* = 0.032) relative to model expectations.

Lines indicate fitted negative binomial model predictions with 95% confidence intervals. Spore density was modeled as a log_10_ transformed predictor. Predictions are shown on the original scale (spores cm^-2^) and plotted on a log_10_ axis. Normal canker count increased nonlinearly with increased spore density with the highest increase occurring between low and moderate densities before plateauing. Susceptible families had consistently higher numbers of normal cankers.

### Dose-response of needle spot severity

Whitebark pine needle spot count increased with spore density but followed a nonlinear response. The quadratic model provided an improved fit compared to the linear model (Table 5). The negative quadratic coefficient suggested a turning point at about 42,0000 spores cm^-2^, which was outside of the highest density tested in the six treatments (max spore density = 24,000 spores cm^-2^). This indicates that spot count had not fully saturated within the experimental inoculum range tested (Fig. 6).

**Figure 6:**
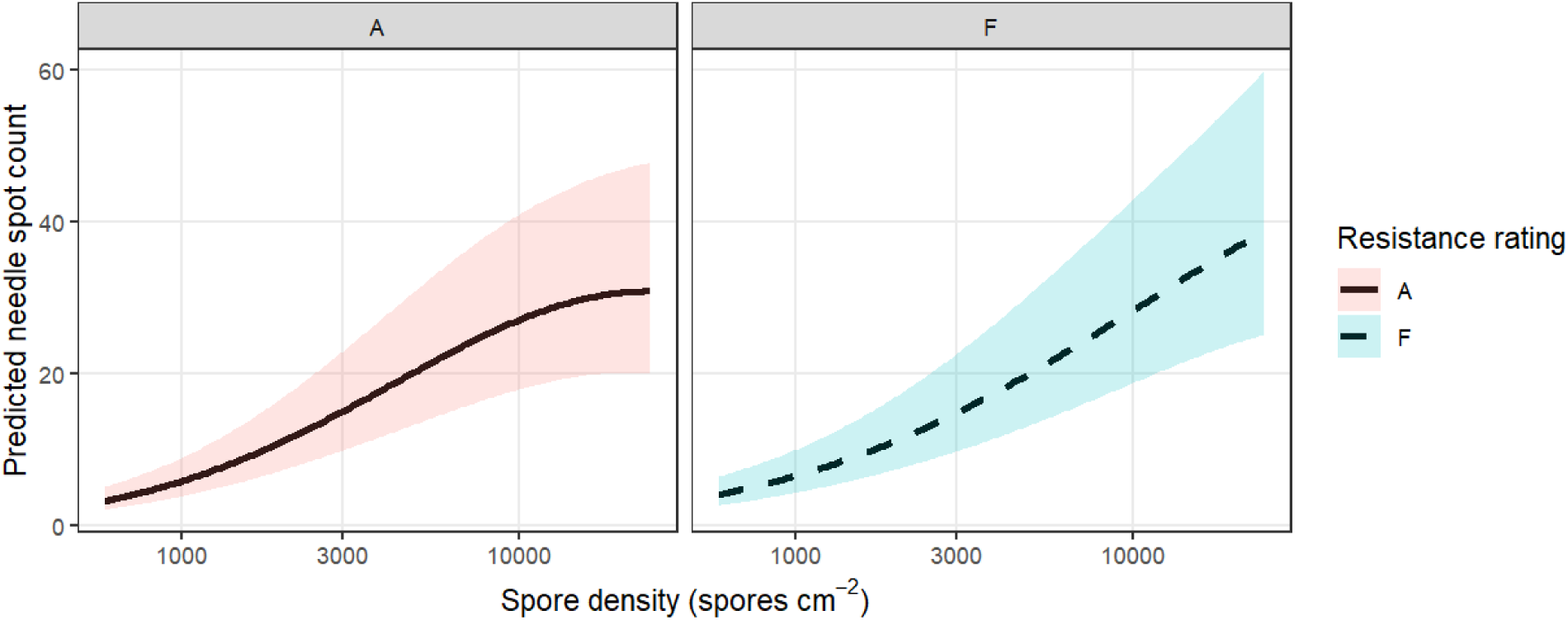
Predicted whitebark pine mean needle spot count across spore densities (spores cm^-2^) for both resistant (A) and susceptible (F) families. Curves are based on negative binomial mixed effects models with family as a random effect and log_10_ transformed spore density modeled as a continuous predictor. Predictions were back transformed to the original scale (spores cm^-2^) and plotted on a log_10_ axis. Needle spot count increased nonlinearly with spore density, with similar scaling observed for both resistance classes. Shaded regions are the 95% confidence intervals.

**Table 5.** Model comparison testing linear vs non-linear dose relationships between spore density and whitebark pine needle spot count. Negative binomial mixed effects models were fitted with seedling family as a random effect and log_10_ transformed spore density modeled as a continuous predictor. The quadratic model significantly improved the fit relative to the linear model (likelihood ratio test χ² = 28.5, Δdf = 1, *p* < 0.001), indicating a nonlinear increase in spot counts with increasing spore density.

| Model | Fixed effects | df | AIC | BIC | Log-likelihood | $\chi^2$ | $\Delta df$ | p-value |
| --- | --- | --- | --- | --- | --- | --- | --- | --- |
| Linear | $\log_{10}(\text{Sdensity})$ | 4 | 4965 | 4983 | -2478.6 | — | — | — |
| Quadratic | $\log_{10}(\text{Sdensity}) + \log_{10}(\text{Sdensity})^2$ | 5 | 4939 | 4961 | -2464.3 | 28.5 | 1 | <0.001 |

Moreover, there was no evidence that resistance rating changed the slope or curvature of the response (interaction *p =* 0.30). This means that both resistant and susceptible families had similar nonlinear scaling of needle spot occurrence with higher inoculum density. This is not unexpected as infection success should saturate because of host leaf area limitation.

### Dose-response of survival

The aggregated probability of survival declined with increased spore density, and the strength of this relationship increased over time (dpi). However, the relationship, in this case, was linear in contrast to spot count for log inoculum density (AIC Linear = 2201.2, Quadratic = 2204.2), which suggests that survival probability declines proportionally with inoculum density (β =-0.66 ± 0.06 SE, *p <* 0.001).

Generally, while needle spot production became limited at high spore density, the risk of mortality continued to scale with pathogen load. A ten-fold increase in spore density resulted in reduced odds of survival by about 69% (β =-0.67 ± 0.06 SE, *p* <0.001). There was no evidence that resistance rating impacted the strength of the response (interaction = 0.93), indicating that both resistant and susceptible families had similar proportional declines in survival with higher spore density.

However, the level of resistance did influence overall survival probability. At the end of the experiment (1,959 DPI), predicted survival differed greatly between resistance classes across the treatments (Fig. 7). Resistant families had much higher survival than susceptible families at both low (71% vs 5%) and high (27% vs <1%) spore densities (Table 6). Even though survival decreased with increasing spore density in both classes, the strength of the dose response was very similar, indicating that QDR mostly shifted overall survival rather than altering sensitivity to increasing rust hazard. In combination, these results indicate that even though needle spot production was constrained at high spore densities, survival probability still scaled with increasing exposure to rust hazard.

**Figure 7.**
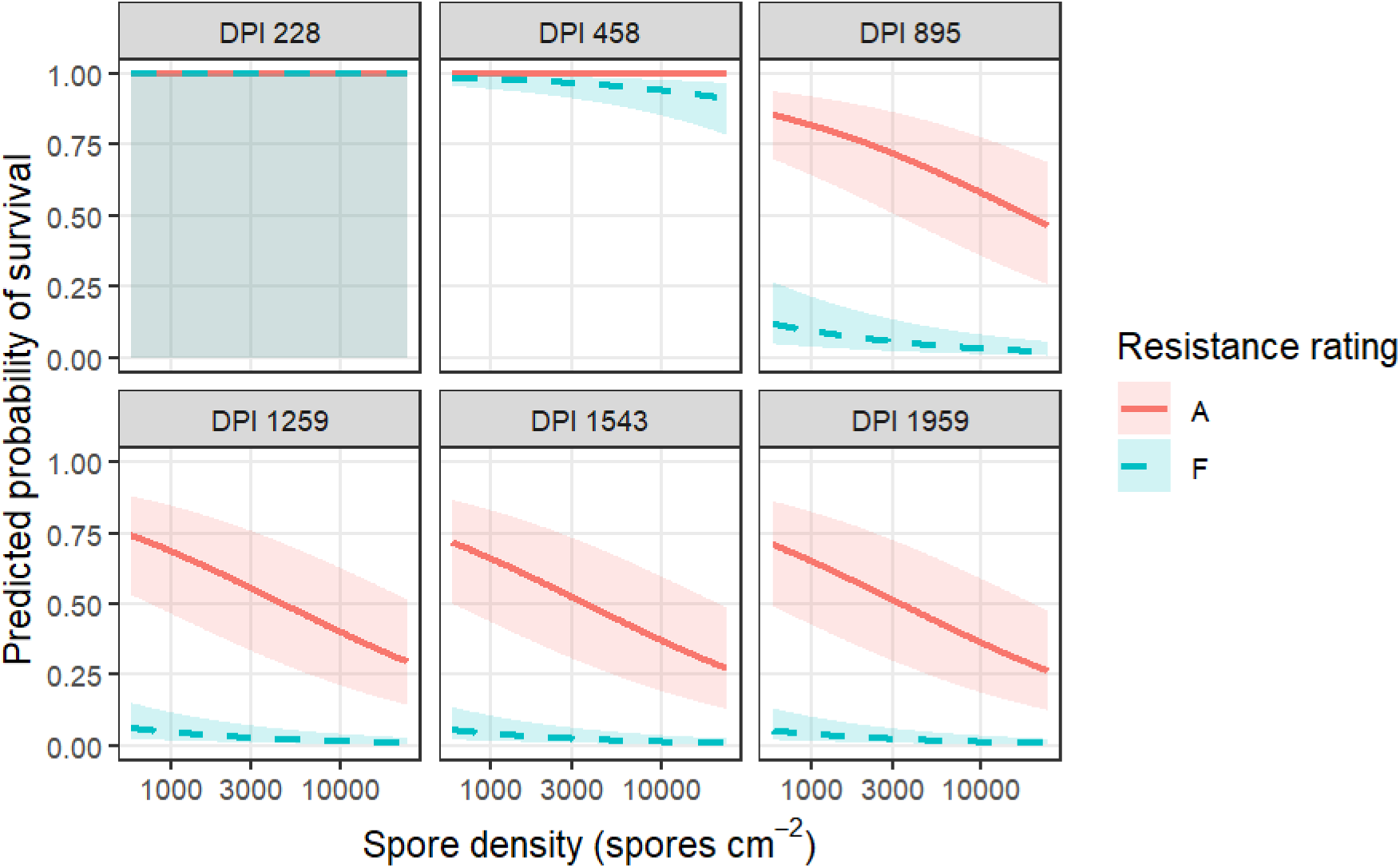
Predicted survival probability across spore density gradients for both resistant (A red) and susceptible (F blue) whitebark pine families at successive inspections (days post inoculation, dpi). Curves are based on binomial mixed effects models with log_10_ spore density and dpi as fixed effects and family as a random effect. Predictions were back transformed and shown on the original scale (spores cm^-2^) and plotted on a log_10_ axis. Survival declined with increased spore density and over time, with resistant families maintaining a much higher survival probability across all densities. Shaded areas represent the 95% confidence intervals.

**Table 6.** Model predicted whitebark pine survival for resistant and susceptible families at the lowest (593 spores cm^-2^) and highest (24,500 cm^-2^) spore densities. Predictions are based on binomial mixed effects models with log_10_ spore density and days post inoculation (dpi) as fixed effects and family as a random effect.

| **Resistance**<br>**Rating** | **Spore**<br>**Density cm<sup>-2</sup>** | **Predicted %**<br>**Survival** |
| --- | --- | --- |
| Resistant | 593 | 71.10% |
| Resistant | 24,500 | 26.60% |
| Susceptible | 593 | 5.30% |
| Susceptible | 24,500 | 0.85% |

### Resistant family differences in survival

Among those families previously rated as resistant (A), survival probability declined with increasing spore density across dpi (Fig. 8). At 228 and 458 dpi, predicted survival remained near 100% across the range of spore densities, suggesting minimal early mortality regardless of spore density. As dpi increased, dose dependent reductions in survival became stronger. By 895 dpi, and at later inspections, survival probability declined steadily with higher spore density, with the steepest declines occurring at the highest densities.

**Figure 8.**
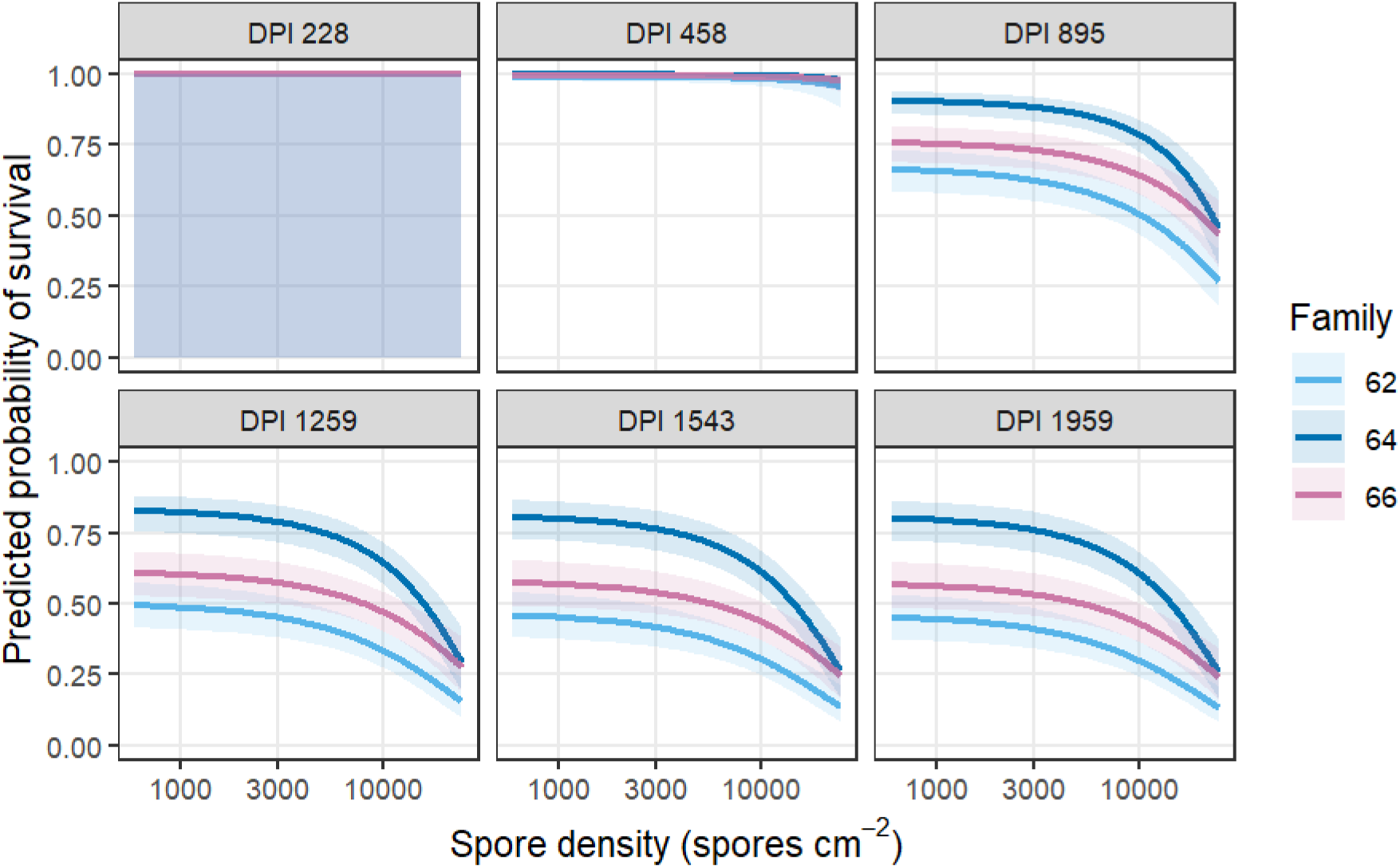
Predicted survival probability across a spore density gradient for individual whitebark pine families rated as resistant (A) at successive days post inoculation (dpi). Curves are based on binomial mixed effects models with log_10_ spore density and dpi as fixed effects and family as a random effect. Spore density was modeled as a log_10_ transformed predictor. Predictions were back transformed and are shown on the original scale (spores cm^-2^) and plotted on a log_10_ axis. Survival declined with increased spore density and over time for all families. Variation in baseline survival was clear, reflecting quantitative disease resistance (QDR) expression. Shaded areas represent the 95% confidence intervals.

Resistant families did differ a little in their overall survival probability, as expected with QDR, but their dose response patterns were mostly the same. For instance, family 64 was consistently more resistant and had higher survival across densities and dpi, while family 62 followed the same pattern but had slightly lower survival. At the highest spore densities, family 64 began to lose its relative resistance advantage and was much closer to family 66 in its overall survival probability.

Model comparisons provided weak support for family-specific slopes of spore density response in survival. The random-slope binomial mixed model, permitting the effect of scaled spore density slope to vary by family, resulted in only marginal improvement relative to the random intercept model (ΔAIC = 0.49). The likelihood ratio test comparing the models was not significant (*X*^2^ = 4.49, df = 1, *p* =0.11), with limited support that resistant families differed in the rate at which survival declined with increasing spore density. QDR leads to differences in baseline survival when exposed to white pine blister rust, but survival probability does not decrease differently given the level of resistance with higher spore densities.

## Discussion

*Cronartium ribicola* has expanded its range over time, and some studies suggest WPBR may be more problematic in the future for some of the high elevation white pine species such as whitebark pine (Dudney et al. 2021, Malone et al. 2026). If this also leads to high to extreme WPBR hazard in some environments, then knowledge of the efficacy of QDR under high spore levels will help forecast the future of the species at those sites. Moreover, very few studies in tree pathosystems have explicitly tested how resistance holds up across a gradient of increasing inoculum density.

In this study, artificial inoculation of whitebark pine seedlings with six levels of increasing *Cronartium ribicola* basidiospore density and known levels of family QDR and susceptibility demonstrated an unmistakably clear and biologically consistent pattern. Increasing spore density dramatically increased infection intensity and mortality risk across all families tested. While the three resistant families retained a strong advantage relative to the susceptible families, that advantage was reduced under heightened disease pressure. These results provide some of the first direct evidence that the effectiveness of QDR in whitebark pine is context dependent, declining as pathogen pressure increases. We are, however, cautiously optimistic that despite the decrease in QDR effectiveness, the probability of survival of resistant trees remains robust and usable even under what looks like extreme exposure.

We found that resistant families had substantially higher survival than susceptible families across all spore densities. Three of the six families in this trial were also in two trials of whitebark pine reported by Sniezko et al. 2024. In those earlier trials, the susceptible (F) families had no survival; while survival of the A rated families exceeded the F families, it was less so in the trial with the slightly higher spore density (trial one spore density 6,013 spores cm^-2^ and trial two 3,359 spores cm^-2^ Sniezko et al. 2024 table 1). Here we found that while resistant families had a survival advantage, the overall magnitude of the advantage declined at higher spore densities.

At low to moderate densities, including those similar to levels commonly used in operational resistance screening (∼3000-4,000 spores cm^-2^) at the USDA Forest Service Dorena Genetic Resource Center (DGRC, Cottage Grove, OR), resistant families maintained large survival differences relative to susceptible families. In seedling inoculation trials, a wide range of resistance among families is often observed (Sniezko et al. 2024). Data from this spore density study indicates that the most susceptible seedling families had little or no survival (99.2% mortality at 24,500 spores cm^-2^), while resistant families had 73.4% mortality. This is a relative reduction in mortality of 26%. The highest spore density used in this study was more than 4000% higher relative to the lowest density and still more than 500% higher than the normal spore density used in operational screening trials at DGRC.

The dose response patterns we observed for both infection and, ultimately, mortality helps to clarify the biological mechanisms that partially underly the expression of QDR. Needle spot counts increased with increasing spore density but did so in a non-linear way, suggesting partial saturation of infection success at high spore densities, most likely reflecting limits imposed by host needle tissue area or other spatial constraints of infection. In contrast, mortality increased linearly with increasing log spore density. This indicates that the cumulative rust burden continues to translate into a higher risk of fatal stem infection as tree defenses become overwhelmed. These findings support a model in which early infection becomes constrained at high spore levels, while downstream progression of the disease (and mortality) is strongly dose dependent in QDR trees.

Field trials have been established to examine performance under natural conditions and validate results from the seedling inoculation trials, but more time is needed for infection levels in the field to be adequate to examine their relationship. The level of spore density in the field is unknown, and trees are exposed for a longer period of time; however, data from a range of white pine species suggests that resistance in the field may be as high or higher than in the operational seedling inoculation trials (Kolpak et al. 2008, Sniezko et al. 2020, Reid et al. 2024).

Seedling inoculation trials are routinely used to evaluate WPBR disease resistance of parent trees. Despite its frequent use, there have been few reports of the effect of increasing inoculum density on effectiveness of QDR, particularly in forest tree species, where most programs are still at an early stage. One case involving *Pinus elliottii* var. *elliottii* Engelm. and its susceptibility to fusiform rust (caused by *C. quercuum* (Berk.) Miyabe ex Shirai f. sp. *fusiforme*) showed that the susceptible family was highly susceptible across all three inoculum densities and two spore sources, while the two resistant families varied in response across these two factors (Griggs et al. 1984). In an agricultural example, increasing inoculum density also interacted with clubroot (*Plasmodiophora brassicae* Woronin) resistant canola (*Brassica napus* L.) where increasing inoculum density had a clear dose-dependent effect (Hwang et al. 2017). In the case of canola, as spore concentration increased, disease severity and gall formation also increased, overcoming resistance at high levels.

The results reported here for whitebark pine demonstrate that at least in some systems spore density can have a large impact on the level of resistance (e.g. survival) reported. However, since few whitebark pine sites in Oregon and Washington appear to have extreme rust hazard levels, it is likely that many of the offspring of resistance families will persist in the presence of WPBR. Field trials currently underway, particularly in areas with known incidence of WPBR, will help monitor the efficacy of resistance.

In the field, disease hazard may vary by environment, and knowing the level of resistance over varying spore densities from screening trials may assist with recommendations on where and how to use QDR. In the trial reported here, the susceptible families showed little or no survival even at the lowest spore density, while survival of the resistant families generally declined somewhat as spore density increased. We suspect that the highest levels of spore density utilized in this trial exceeds what may occur even on the sites with the highest WPBR hazard for whitebark pine, but they provide information for land managers planning restoration activities of whitebark pine. The range of QDR in whitebark pine is generally much higher than that in other white pine species such as sugar pine or western white pine (Kegley and Sniezko 2004), so higher inoculum density may obscure the differences between susceptible and QDR in those species and in other pathosystems where the level of QDR from wild populations is relatively low level. The results also provide impetus to other forest tree resistance programs to investigate the impact of spore density on QDR and to ensure that some field trials are present on sites with high disease hazard.

## Acknowledgments

We thank Kristin Silva and Emily Boes for assistance with some of the assessments, and others at Dorena Genetic Resource Center who assisted with other phases of the trial as well as those in USDA Forest Service and National Park Service that collected the seedlots.

## CRediT Author contributions

Jeremy S. Johnson: Conceptualization, Methodology, Investigation, Formal analysis, Visualization, Writing – original draft, Writing – review & editing. Benjamin Wilhite: Methodology, Investigation, Data curation, Formal analysis, Visualization, Writing – original draft, Writing – review & editing. Robert Danchok: Investigation, Writing – review & editing. Angelia Kegley: Methodology, Investigation, Writing – original draft, Writing – review & editing. Richard A. Sniezko: Conceptualization, Methodology, Investigation, Formal analysis, Writing – original draft, Writing – review & editing, Supervision, Project administration.

## References

Government of Canada. (2012) Order amending Schedule 1 to the Species at Risk Act. Canada Gazette Part II 146:SOR/2012-2113.

U.S. Fish and Wildlife Service. (2022) Endangered and threatened wildlife and plants; Threatened species status with section 4(d) rule for whitebark pine (*Pinus albicaulis*). Federal Register 87 (240): 76882–76917.

Dudney, J., C. E. Willing, A. J. Das, A. M. Latimer, J. C. B. Nesmith, and J. J. Battles. 2021. Nonlinear shifts in infectious rust disease due to climate change. Nature Communications 12:5102.

Geils, B. W., K. E. Hummer, and R. S. Hunt. 2010. White pines, ribes, and blister rust: a review and synthesis. Forest Pathology 40:147–185.

Griggs, M. M., R. J. Dinus, and G. A. Snow. 1984. Inoculum source and density influence assessment of fusiform rust resistance in slash pine. Plant Disease 68:770–774.

Hartig, F. 2026. DHARMa: Residual diagnostics for hierarchical (multi-level/mixed) regression models. R package.

Hoff, R. J. 1986. Inheritance of the bark reaction mechanism in *Pinus monticola* infected by *Cronartium ribicola*. USDA Forest Service, Intermount Research Station, Research Note INT-361.

Hoff, R. J. 1992. How to recognize blister rust infection on whitebark pine. USDA Forest Service Intermountain Research Station Research Note INT-406.

Hoff, R. J., R. T. Bingham, and G. I. McDonald. 1980. Relative blister rust resistance of white pines. European Journal of Forest Pathology 10:307–316.

Hoff, R. J., D. Ferguson, G. I. McDonald, and R. E. Keane. 2001. Strategies for managing whitebark pine in the presence of white pine blister rust Pages 346–366 *in*D. Tomback, S. F. Arno, and R. E. Keane, editors. Whitebark pine communities: Ecology and restoration. Island Press, Washington.

Hoff, R. J., and G. McDonald. 1980. Improving rust-resistant strains of inland western white pines. USDA For. Serv. Res. Pap. INT-245, 13 p. Intermt. For. and Range Exp. Stn., Ogden, Utah.

Hwang, S. F., S. E. Strelkov, H. U. Ahmed, V. P. Manolii, Q. Zhou, H. Fu, G. Turnbull, R. Fredua-Agyeman, and D. Feindel. 2017. Virulence and inoculum density-dependent interactions between clubroot resistant canola (Brassica napus) and Plasmodiophora brassicae. Plant Pathology 66:1318–1328.

Johnson, J. S., and R. A. Sniezko. 2021. Quantitative disease resistance to white pine blister rust at wouthwestern white pine’s (*Pinus strobiformis*) northern range. Frontiers in Forests and Global Change 4.

Kegley, A. J., and R. A. Sniezko. 2004. Variation in blister rust resistance among 226 *Pinus monticola* and 217 P. *lambertiana* seedling families in the Pacific Northwest. Pages 209-225 *in* Breeding and genetic resources of five needle pines: Genetics, breeding and adaptability, Proceedings of the IUFRO 2.02.15 Working Party Conference USDA Forest Service, Rocky Mountain Research Station RMPS-P-32. Fort Collins, CO., Medford, OR.

Kinloch, B. B. 2003. White pine blister rust in North America: Past and prognosis. Phytopathology 93:1044–1047.

Kinloch, B. B., and G. E. Dupper. 2002. Genetic specificity in the white pine-blister rust pathosystem. Phytopathology 92:278–280.

Kinloch, B. B., G. K. Parks, and C. W. Fowler. 1970. White pine blister rust: Simply inherited resistance in sugar pine. Science 167:193–195.

Koester, H., D. P. Savin, M. Buss, and R. A. Sniezko. 2018. White pine blister rust hazard rating for 265 sites in southern Oregon, USA. Pages 173-180 *in* Genetics of five-needle pines, rusts of forest trees, and Strobusphere. Rocky Mountain Research Station, Fort Collins, CO.

Kolpak, S. E., R. A. Sniezko, and A. J. Kegley. 2008. Rust infection and survival of 49 *pinus monticola* families at a field site six years after planting. Annals of Forest Research 51:67–80.

Liu, J. J., J. S. Johnson, and R. A. Sniezko. 2021. Genomic advances in research on genetic resistance to white pine blister rust In in North American white pines.*in* A. R. De La Torre, editor. The Pine Genomes. Springer Nature, Switzerland.

Mahalovich, M. F., and L. Stritch. 2013. Pinus albicaulis. The IUCN Red List of Threatened Species: e.T39049A2885918. 10.2305/IUCN.UK.2013-1.RLTS.T39049A2885918.en.

Malone, S. L., A. W. Schoettle, K. S. Burns, H. S. J. Kearns, J. E. Stewart, M. Newcomb, and C. M. Cleaver. 2026. Future climate will not save high-elevation white pines. Communications Earth & Environment.

RCoreTeam. 2026. R: a language and environment for statistical computing. R Foundation for Statistical Computing, Vienna, Austria.

Reid, I. R., C. Cartwright, R. A. Sniezko, R. C. Hamelin, and S. N. Aitken. 2024. Field-testing whitebark pine resistance to white pine blister rust: A simple, effective approach to progeny testing for restoration. Forest Ecology and Management 553:121647.

Schwandt, J. W. 2006. Whitebark pine in peril: a case for restoration. U.S. Department of Agriculture, Forest Service, Northern Region, Forest Health Protection, Missoula, MT.

Sniezko, R. A., J. S. Johnson, A. Kegley, and R. Danchok. 2024. Disease resistance in whitebark pine and potential for restoration of a threatened species. Plants, People, Planet 6:341–361.

Sniezko, R. A., J. S. Johnson, and D. P. Savin. 2020. Assessing the durability, stability, and usability of genetic resistance to a non-native fungal pathogen in two pine species. Plants, People, Planet 2:57–68.

Sniezko, R. A., A. Kegley, and R. Danchok. 2008. White pine blister rust resistance in North American, Asian and European species - results from artificial inoculation trials in Oregon. Annals of Forest Research 51:53–66.

Sniezko, R. A., A. J. Kegley, R. Danchok, and S. Long. 2007. Variation in resistance to white pine blister rust among 43 whitebark pine families from Oregon and Washington—early results and implications for conservation. Pages 82–97 *in* Whitebark pine: a Pacific Coast perspective. U.S. Department of Agriculture, Forest Service, Pacific Northwest Region, Ashland, OR.

Sniezko, R. A., and J.-J. Liu. 2022. Genetic resistance to white pine blister rust, restoration options, and potential use of biotechnology. Forest Ecology and Management 520:120168.

Sniezko, R. A., and J. J. Liu. 2023. Prospects for developing durable resistance in populations of forest trees. New Forests 54:751–767.

Sniezko, R. A., M. F. Mahalovich, A. W. Schoettle, and D. R. Vogler. 2011. Past and current investigations of the genetic resistance to *Cronartium ribicola* in high-elevation five-needle pines. Pages 246-264 *in* R. E. Keane, D. F. Tomback, M. P. Murray, and C. M. Smith, editors. The future of hight-elevation, five-needle white pines in western North America. Proc High Five Symp RMRS-P-63, USDA Forest Service Rocky Mountain research Station, Fort Collins, CO.

Sniezko, R. A., J. Smith, J.-J. Liu, and R. Hamelin. 2014. Genetic resistance to fusiform rust in southern pines and white pine blister rust in white pines—A contrasting tale of two rust pathosystems—Current status and future prospects. Forests 5:2050–2083.

Tomback, D. F., and P. Achuff. 2010. Blister rust and western forest biodiversity: ecology, values and outlook for white pines. Forest Pathology 40:186–225.

Tomback, D. F., R. E. Keane, A. W. Schoettle, R. A. Sniezko, M. B. Jenkins, C. R. Nelson, A. D. Bower, C. R. DeMastus, E. Guiberson, J. Krakowski, M. P. Murray, E. R. Pansing, and J. Shamhart. 2022. Tamm review: Current and recommended management practices for the restoration of whitebark pine (Pinus albicaulis Engelm.), a threatened high-elevation Western North American forest tree. Forest Ecology and Management 522:119929.

Tomback, D. F., and E. Sprague. 2022. The National Whitebark Pine Restoration Plan: Restoration model for the high elevation five-needle white pines. Forest Ecology and Management 521:120204.

